# Foot placement control underlies stable locomotion across species

**DOI:** 10.1101/2024.09.10.612345

**Authors:** Antoine De Comite, Nidhi Seethapathi

## Abstract

Animals navigate their environment stably without inefficient course corrections despite unavoidable errors. In humans, this stability is achieved by varying the placement of the foot on each step such that recent movement errors are corrected. However, it is unknown how animals with diverse nervous systems and body mechanics use foot placement control: foot trajectories of many-legged animals are thought to be stereotypical velocity-driven patterns, as opposed to error-driven. Here, we posit a unified “feedforward-feedback” control structure for foot placement by combining velocity-driven and body state error-driven contributions. We provide empirical support for this control structure across flies, mice and humans by mining the variability in the foot placements and body states during natural locomotion. We find that a competing “feedforward-only” control structure with purely velocity-driven foot placement is not supported by the data. This work discovers shared behavioral signatures of foot placement control in flies, mice, and humans. The urgency and centralization of these control signatures vary with the animal’s neuromechanical embodiment; more inherently stable many-legged embodiment is associated with a lower control magnitude and timescale. Further, many-legged embodiment is accompanied by modular direction- and leg-specific signatures, which are centralized across both legs in humans. Taken together, our findings provide insight into stabilizing foot placement control across species, revealing how different neuromechanical embodiments achieve a shared functional goal.

## Introduction

Stable locomotion is a fundamental motor skill for legged species as it enables navigating the environment without time-consuming course corrections in the presence of intrinsic sensorimotor noise or external perturbations. However, we lack a unified understanding of how this important function is achieved in species with diverse neural control and body mechanics, i.e. across *neuromechanical embodiment*. In human and robot locomotion, stability is achieved through foot placement control [1, 2], where the swinging leg is placed in a manner that corrects recent deviations in the body’s state [3, 4, 5]. In contrast, foot placements in multi-legged animals are predominantly thought to be top-down and velocity-dependent [6, 7], hindering comparative insight into error-dependent locomotor control. Here, we hypothesize that a unified control structure underlies stable locomotion across legged species, combining velocity- and error-dependent components, and develop a data-driven model to identify control signatures from natural locomotion across species. We find that flies (*Drosophila melanogaster*), mice, and humans exhibit shared signatures of velocity- and body state error-dependent foot placement control. These findings enable comparative study of the neuromechanics and motor control of locomotor stability.

Understanding the neuromechanical basis of motor control requires comparative insight into how different species achieve the same behaviorally-relevant functional goal [8, 9]. While the behavioral signatures of stereotypical velocity-dependent foot placement patterns across species [10, 11, 12, 13, 14, 15, 16] has helped solidify the existence of velocity-driven neural control during locomotion [17, 6], we lack analogous cross-species signatures of error-driven foot placement. Indeed, while there is agreement about error-based foot placement control in humans [3, 18, 4], we lack comparable evidence in other legged animals. Such error-based foot placement control could rely on diverse sensory feedback [19, 20, 21, 22, 23, 24] and neural processing [25, 21, 26, 27], and its neural underpinnings are challenging to establish in humans. Identifying common signatures of stabilizing foot placement control across species is an important step towards obtaining a deeper understanding of its functionally-relevant neuromechanical basis. To address this gap we develop a data-driven model to identify signatures of foot placement control across species. Using this model, we discover that mice and flies exhibit human-like signatures of stabilizing foot placement control.

Explaining how animals achieve stable locomotion has been a long-standing target for theoretical modeling. Two primary forms of theoretical control structure hypotheses have emerged to explain how animals walk and run stably. The first one posits that a propulsive velocity-increasing action alone is sufficient for stability in the presence of self-stabilizing passive mechanics [28, 29, 30, 31]. In this hypothesized control structure, a “feedforward” controller prescribing the swing leg kinematics and contact timing purely as a function of movement velocity [32, 33] is sufficient to guarantee stability [34, 35]. Other theoretical models suggest that such feedforward velocity-dependent strategies may be insufficient to guarantee stability in the lateral plane [36]. Thus, a second control structure hypothesis posits a “feedback” controller which needs to measure past errors to dictate current foot placement [36, 37, 38]. Yet, such velocity-dependent and error-dependent control has not, to the best of our knowledge, been unified into a single control structure hypothesis as we posit here. Moroeover, although both feedforward and feedback control structures have been shown theoretically to to enable stabilization [34, 35, 36], there is a lack of empirical evidence. To address this gap, we first extend the feedforward velocity-dependent control hypothesis to lateral foot placement, then define a unified control structure with body state error-based control in the of a velocity-dependent module, and finally identify data-driven signatures to test these hypotheses.

An animal’s neuromechanical embodiment, i.e. interactions between the mechanical properties of the body and the neural control strategies, influences its locomotion [39, 40]. Biomechanical factors such as passive muscle mechanics [41], body size-dependent transmission delays [42, 43], or the configuration of the legs [44, 45] can shape the animal’s chosen control strategies. Neural control of locomotion is achieved through spinal circuits [6, 46] and is affected by perturbations to different neural centers [47, 48, 49, 50, 51]. Even though the influence of these neuromechanical factors on feedforward velocity-dependent locomotion is well-established [39, 52, 7, 53, 41], how they affect stabilizing error-dependent control remains poorly understood. Animals with diverse neuromechanical embodiments may adjust their foot placement control strategies differently in response to errors, but we lack a cross-species characterization of foot placement control needed to study such variations. Here, we put forth a cross-species generalizable data-driven model identifying error-dependent signatures of foot placement control, highlighting how they change with multi-legged neuromechanical embodiment.

In this study, we investigate stabilizing foot placement control across species with diverse neuromechanical embodiment, discovering signatures of a feedforward-feedback control structure across species. Our approach and findings have implications for comparative neuromechanics, comparative neuroscience, and legged robotics.

## Results

We hypothesize that a similar feedforward-feedback control structure underlies stabilizing foot placement in any legged animal, and discovered signatures of this control structure hidden in natural locomotor variability. We characterized these signatures in flies (*Drosophila melanogaster*), mice (C57BL/6J), and humans using high-throughput locomotion data, discovering shared velocity-dependent and body state error-dependent signatures across species, and highlighting how the signatures change with more inherently stable and multi-legged embodiment.

### Feedforward-only and feedforward-feedback control structure hypotheses for foot placement

We evaluate two hypothesized control structures underlying foot placement across species. We use the terms “feedforward” and “feedback” to refer to the theorized control structures such as using forward velocity alone as an input [54] or also using body state error as an input [37]. Note that we do not intend these terms to refer to the neural basis of the control.

The first control structure is feedforward-only (Figure 1A, black box), transforming velocity and heading inputs into nominal velocity-dependent foot placement patterns for each species, reflecting their characteristic gait patterns (Figure 1B). We rationalize this controller structure with animal neuroscience research, which suggests that velocity and heading are important descending inputs for the generation of periodic inter-limb coordination patterns [55, 56, 57]. For this feedforward-only control structure, we generalize the leg length and hip angle actions in prior theoretical work [54] to foot placement location and timing in our space of observable variables. Moreover, we generalize these prior theoretical works to lateral feedforward-only stabilizing control, by hypothesizing that the velocity-dependent changes in the animals lateral foot placement location and timing could have a stabilizing effect (Figure 1E), given the variability in the lateral swing foot kinematics both as a function of gait phase and forward velocity.

**Figure 1:**
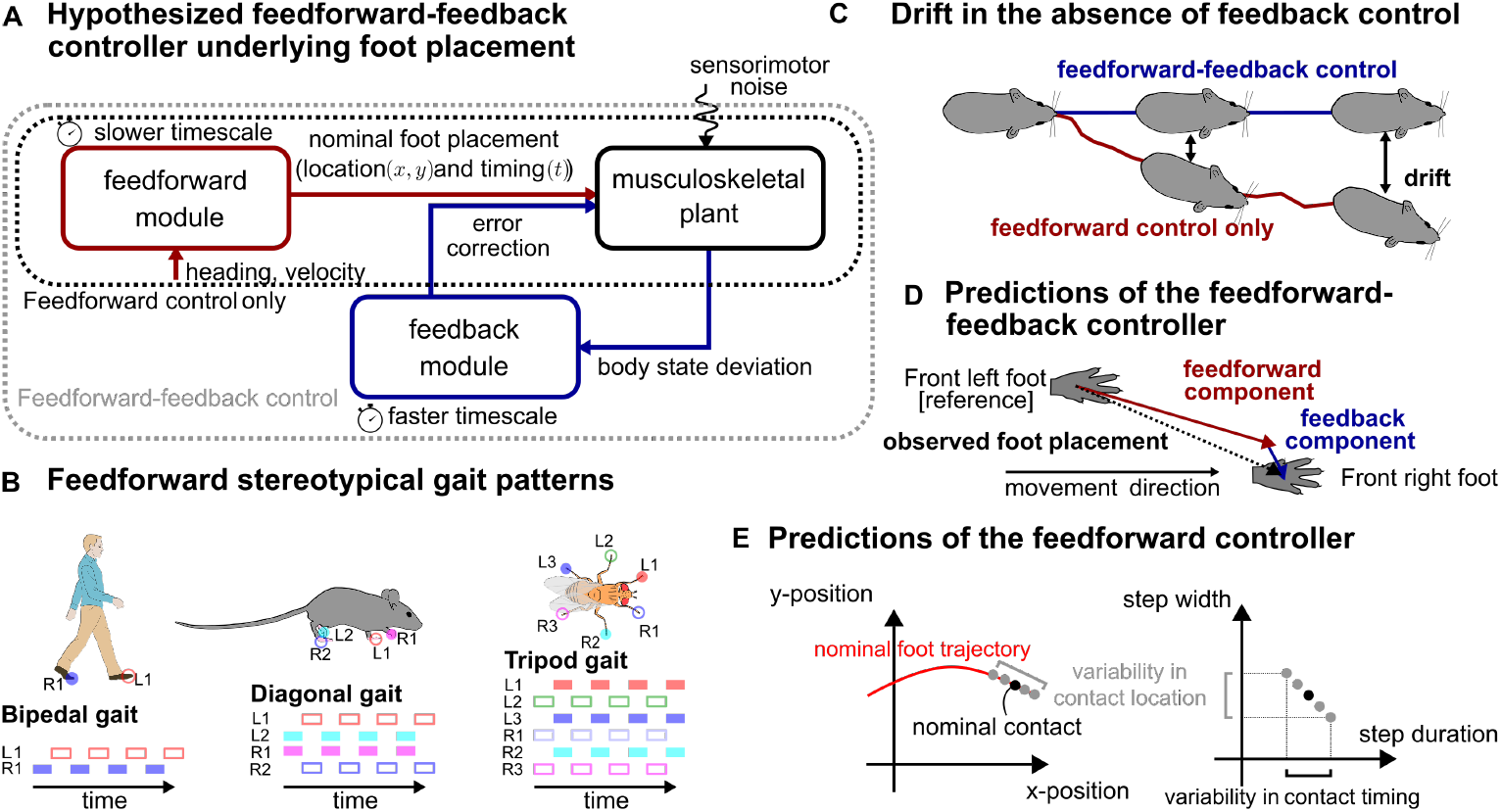
Hypothesized control structure for stable locomotion in legged animals. **A** Schematic representation of the two alternative control structures underlying foot placement as well as the interactions between the velocity-dependent feedforward control module (red) and the body state error-dependent feedback control module (blue). The feedforward module transforms velocity and heading inputs into nominal foot placements over multiple gait cycles. The feedback module corrects body state errors by making appropriate adjustments to the foot placement. **B** Canonical gait patterns for humans, mice and flies. Colored rectangles indicate the times at which each limb is in stance, full and empty rectangles illustrate the two groups of legs alternatively in contact with the floor during the specific gait patterns studied. **C** Positional drift due to intrinsic sensorimotor noise in presence (blue) and absence (red) of body state error-based feedback control. **D** Under the feedforward-feedback assumption, the observed foot placement is the sum of the individual contributions of feedforward and feedback modules. **E** The velocity-dependent nominal trajectory of the foot (red line, left) prescribes the position of the foot as a function of time. A slightly shorter or longer contact timing could results in deviations of the contact location from its nominal position (gray dots vs black dot). Therefore, the variability in contact timing could result in apparent variability in contact location even in absence of body state error-based information.

The second hypothesized control structure is hierarchical and feedforward-feedback, consisting of two different control modules (Figure 1A, gray box): a velocity-dependent module and a body state error-dependent module. The body state error-dependent feedback module is defined in the vicinity of the feedforward module, to correct deviations from the intended velocity-dependent nominal trajectory. This feedforward-feedback controller is hierarchical as each module has access to distinct inputs and operates at different timescales. The feedforward module transforms the velocity and heading inputs into nominal foot placement patterns over a timescale of multiple gait cycles, while the feedback control module transforms the body state errors, i.e. deviations between the actual body state and the nominal behavior, into corrective foot placements at a faster timescale (Figure 1A). Importantly, these body state errors are defined as deviations from the velocity-dependent nominal behavior and not from the average behavior as done in previous work [3, 58, 59], which allows us to unify error-dependent and velocity-dependent foot placement control. We rationalize the body-state error-dependent feedback module with human locomotor stability research on body state error-based modulation of foot placement in healthy [18, 3] and disordered locomotion [60, 61, 27, 62, 26]. Under the assumption of this control structure, the net observed foot placement would be the resultant of contributions from each control module (Figure 1D).

If the feedforward-only controller is sufficient to explain foot placement, we should find that the velocity-dependent module explains an equal or greater amount of the variability in the foot placement location and timing than the body state error-dependent module. Alternatively, if the feedforward-feedback control structure underlies foot placement control, we should find that the body state error-dependent module explains a significant amount of the variability in foot placement location and timing that is not captured by the velocity-dependent module alone (Figure 1D).

### Signatures of velocity-dependent feedforward foot placement across species

The feedforward control module (Figure 1A, red) posits that movement velocity contributes to the foot placement patterns adopted by animals. Although the specific contribution of velocity to both fore-aft and lateral foot placement variability has been characterized in humans [63], this relationship is less understood across species. To simultaneously test our hypothesized feedforward module and understand the nature of its velocity-dependence in multi-legged animals, we take a data-driven approach, correlating the movement velocity and the foot placement location and timing variability.

We find a consistent positive correlation between movement velocity and step length across all species and limbs, but not between velocity and step width. In flies, step length for each leg is positively correlated with movement velocity (chatterjee-xi test: *p <* 0.05 for all legs; Figure 2C, left) while step width shows no correlation (chatterjee-xi test: *p* = 0.924 (front), *p* = 0.3445 (middle), and *p* = 0.138 (hind); Figure 2C, center). Post-hoc regressions confirm that flies linearly increase their step length with increasing movement velocity (significant in 8/8 videos for front, central and hind limbs). Mice also show a positive correlation between step length and velocity (chatterjee-xi test: *p <* 0.05 for all legs; Figure 2B, left) but not step widths (chatterjee-xi test: *p* = 0.15 (front) and *p* = 0.0636 (hind), Figure 2B, center). Similar to flies, mice linearly increase their step length with increasing velocity for both front (significant in 79/80 animals) and hind limbs (significant for 44/80 animals). In humans, we observe a similar pattern: step length increase with increasing velocity (chatterjee-xi test: *p <* 0.05, post-hoc significant for 20/21 subjects, Figure 2A, left), while step width remain constant (chatterjee-xi test: *p* = 0.06, Figure 2A, center). Finally, we find a negative correlation between step duration and velocity (chatterjee-xi tests: *p <* 0.05 for all legs and all species, Figure 2A-C, right) and find that step duration exponentially decreases with increasing velocity across all species.

**Figure 2:**
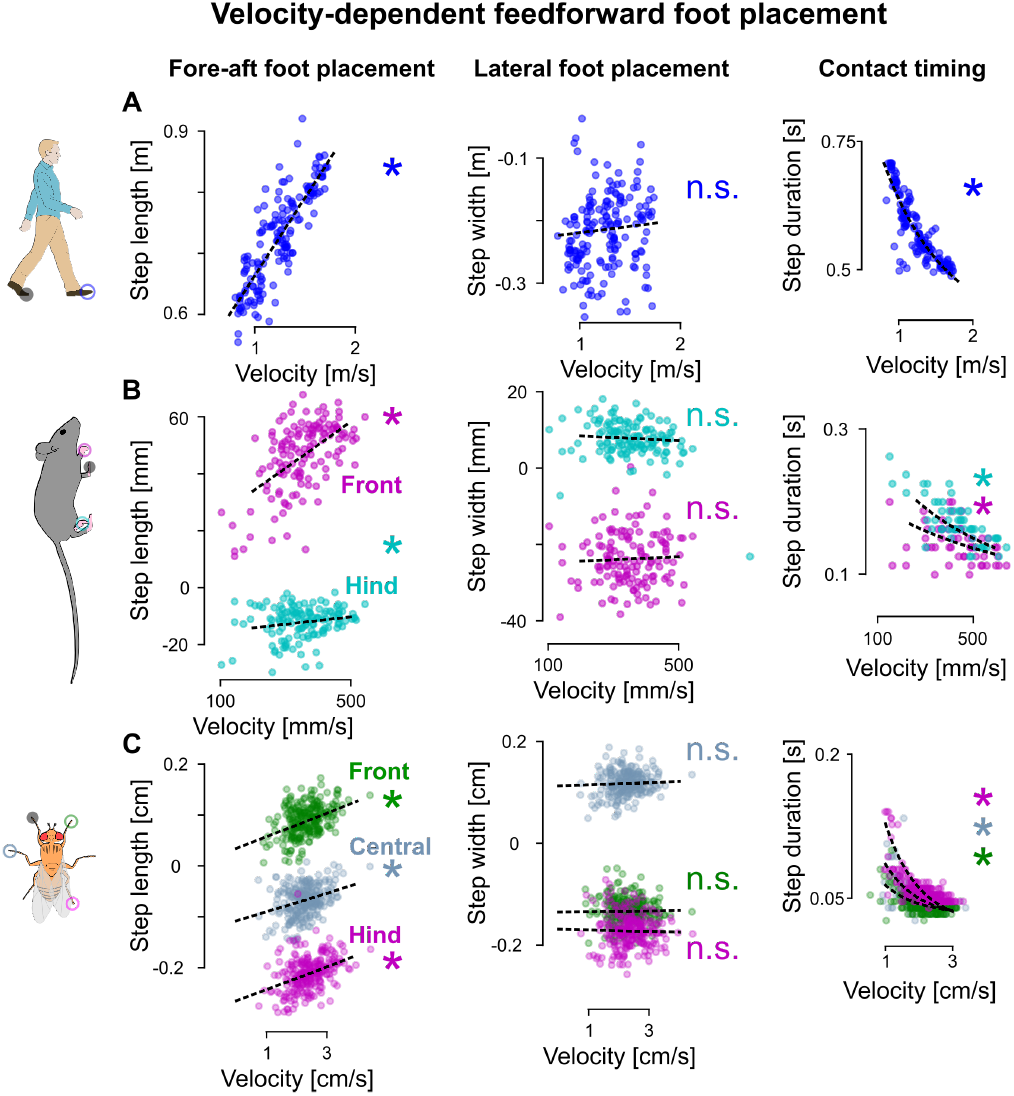
Signatures of velocity-dependent feedforward foot placement control across species. **A** Modulation of step length (left), step width (center), and step duration (right) with movement velocity for humans. **B** Modulation of step length (left), step width (center), and step duration (right) with movement velocity for front (magenta) and hind (cyan) limbs of mice. **C** Modulation of step length (left), step width (center), and step duration (right) with movement velocity for front (green), central (blue), and hind (magenta) limbs of flies. The black dashed lines capture the individual regressions between movement velocity and the step width/length. The reference limbs are represented as black disk overlaid on the animal cartoon, the open disks represent the different feet studied. ∗ : *p <* 0.05

The step width variability that is not explained by movement velocity could stem from variability in contact timing, as suggested by the feedforward-only controller (Figure 1E). To investigate this possibility, we sought potential correlations between the deviations in contact timing from its nominal behavior (Figure 2, right column) and the deviations the step width from its nominal behavior (Figure 2, central column). We did not find such correlations for any of the species (chatterjee-xi test: *p >* 0.05), which strongly suggests that the feedforward-only hypothesis might not be the sole source of observed step width variability.

Thus, we find a similar influence of movement velocity on nominal foot placement across across all species and limbs. Specifically, movement velocity modulates both step length and step duration but not the step width in all species. Foot placement timing variations also do not explain the variance in step width. Faster movement velocities result in longer steps and shorter step durations, while step width remain largely unaffected by the hypothesized feedforward-only control structure.

### Signatures of body state error-dependent feedback foot placement across species

Our hypothesized feedback module (Figure 1A, blue) suggests that errors in the body state on each step should inform the future foot placement. To test this hypothesis and characterize the feedback control module, we map the kinematic body state errors with the future foot placement, quantifying the variance in foot placement explained by this mapping as a function of gait fraction. Such a mapping has previously been identified during human treadmill locomotion [3, 64, 5], but its existence in other legged species and during natural overground human locomotion remains unknown. To characterize this mapping across legged species, we calculate body state and foot placement errors, defined as deviations from the nominal patterns predicted by the velocity-dependent model, at various gait fractions (Figure 3A) and identify linear relationships between the deviations in the body state and the foot placements (see *Methods*). We then compare these mappings to a baseline mapping derived from foot kinematics alone to determine whether body state errors contain more information about future foot placement than the foot kinematics itself.

**Figure 3:**
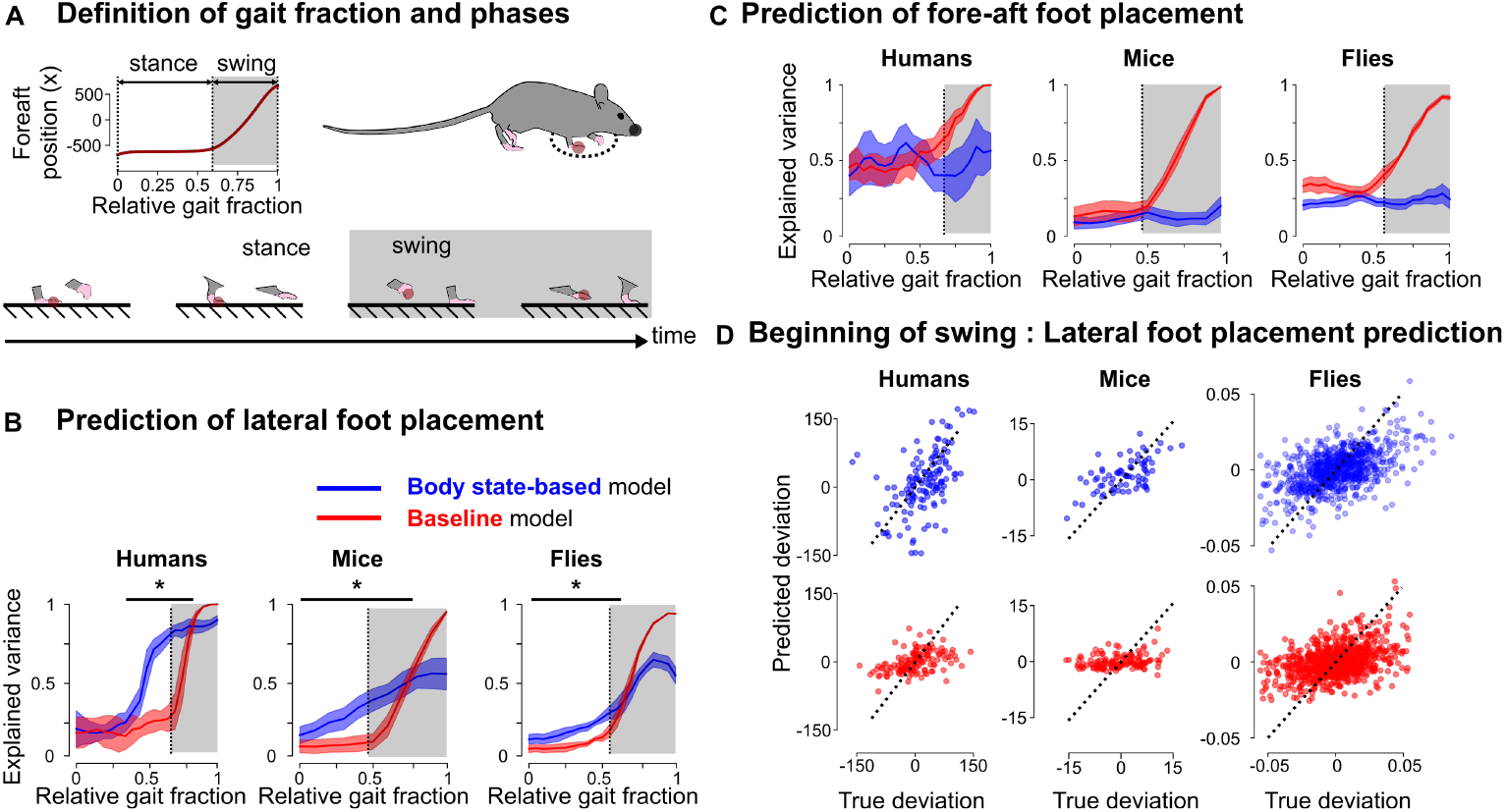
Signatures of body state error-dependent feedback foot placement control. **A** We defined the gait fraction as the time elapsed since the last contact of the reference foot divided by the duration of the gait cycle. The stance and swing phase of each foot (here illustrated for the foot highlighted in red) were defined as the periods during which the foot was or was not in contact with the ground, illustrated here for one of the front limbs in mice. The swing-stance and stance-swing transitions correspond to contact initiation and contact termination. We built models to predict future contact initiation based on the kinematics of the body during the gait cycle preceding that contact. **B** Explained variance (R^2^, median and interquartile range) obtained for the body state-based and baseline predictions of the lateral deviation in foot placement as a function of the relative gait fraction. The gray areas represent the swing phase of the foot to be placed, the vertical dashed lines represent the stance-swing transition for each species. **C** Explained variance (R^2^, median and interquartile range) obtained for the body state-based and baseline prediction predictions of the fore-aft foot placement as a function of the relative gait fraction. The gray areas represent the swing phase of the foot to be placed, the vertical dashed lines represent the stance-swing transition for each species. **D** Comparison between the true and predicted deviations for the body state-based (top, blue) and baseline (bottom, red) predictions of the lateral foot placement of humans, mice and flies front limbs at the beginning of the swing phase of the contact foot (dashed line in panel **B**). The black dashed lines represent the identity lines corresponding to perfect prediction. ∗ : *p <* 0.05

We find that body state errors predict future lateral foot placement in the front limbs of flies, mice, and humans. In all species, these body state error-based predictions outperform the baseline predictions, derived from the foot kinematics, well before swing onset. In flies, the body state-based predictions are superior (Kolmogorov-Smirnov test (*p <* 0.05), Figure 3B, right) from 0% to 65% of the gait cycle, while swing onset occurs around 55%. In mice, the body state error-based predictions are superior (Kolmogorov-Smirnov test (*p <* 0.05), Figure 3B center) from 0% to 70% of the gait cycle, while swing onset occurs around 50%. In humans, the body state error-based predictions are superior (Kolmogorov-Smirnov test (*p <* 0.05), Figure 3B left) from 30% to 80% of the gait cycle, while swing onset occurs around 65%. In all three species, the body state-based model predicts deviations more accurately than foot kinematics based model well before swing onset (Figure 3C, representative data). We also analyze the fore-aft foot placement in a similar manner and find no significant differences across all three species (Figure 3C); the body state error-based model prediction is never better than the baseline model for fore-aft foot placement.

We also analyzed the eigenvalues of the Poincare return maps for all three species. Eigenvalues with a modulus lower than 1 indicate a stable gait. We find that the modulus of all eigenvalues is indeed smaller than 1 for all three species, with the spectral radii (the modulus of the largest eigenvalue) decreasing with increasing number of legs (humans *ρ* = 0.98, mice *ρ* = 0.65, and flies *ρ* = 0.48). To determine whether the body state error-based foot adjustments contribute to stability, we next investigate whether lateral variations in foot placement contribute to error reduction. We reason that if these variations play a stabilizing role, larger foot placement deviations should be associated with larger reductions in body state errors from one half-gait cycle to the next (see *Methods*). We find a significant positive correlation between foot placement deviation and subsequent error reduction in all three species (humans: *p <* 0.05 and *β* = 0.49, mice: *p <* 0.05 and *β* = 0.29, flies: *p <* 0.05 and *β* = 0.87, Figure S5). Notably, this analysis between foot placement deviation and error reduction can be reformulated as *differences-in-differences*, which is a quasi-experimental causal inference approach (See section S3 for details). This finding suggests that the observed mappings between body state error and lateral foot placement can have a stabilizing effect on the body.

Thus, we find that body state errors predict future lateral foot placement in the front limbs of mice and flies, similar to humans. We also provide evidence that these error-based foot placements could have a stabilizing effect, reflected in the finding that past foot placement magnitude is correlated with future reduction in body state errors. These findings are consistent with the body state error-based control structure hypothesis for lateral foot placement during natural locomotion across species.

### Signatures of control magnitude and timescale vary across embodiment

We have found qualitatively similar body state error-dependent lateral foot placement control signatures, hidden in locomotor variability, in flies, mice, and humans. However, the details of the control strategy – such as how much error is corrected and over what timescale [65]– may vary across species due to more passively stable embodiment afforded by multiple limbs. To understand how the amount of error corrected in each step, which we term the *control magnitude*, varies with embodiment we computed the maximal difference between the coefficient of determination of the body state error-based model relative to the foot trajectory-based baseline model (Figure 4A insert) for each individual regression, and compared it across species. We observed a gradient across species (Figure 4A) with the largest differences observed in humans (0.61 ± 0.14), followed by mice (0.31 ± 0.12) and by flies (0.14 ± 0.04). These results suggested that, although these species leveraged qualitatively similar body state error-dependent foot placement signatures, the extent to which errors are corrected in a single step vary with many-legged embodiment. Specifically, the explained variance by body state error-based control decreased with increasing passively stable embodiment.

**Figure 4:**
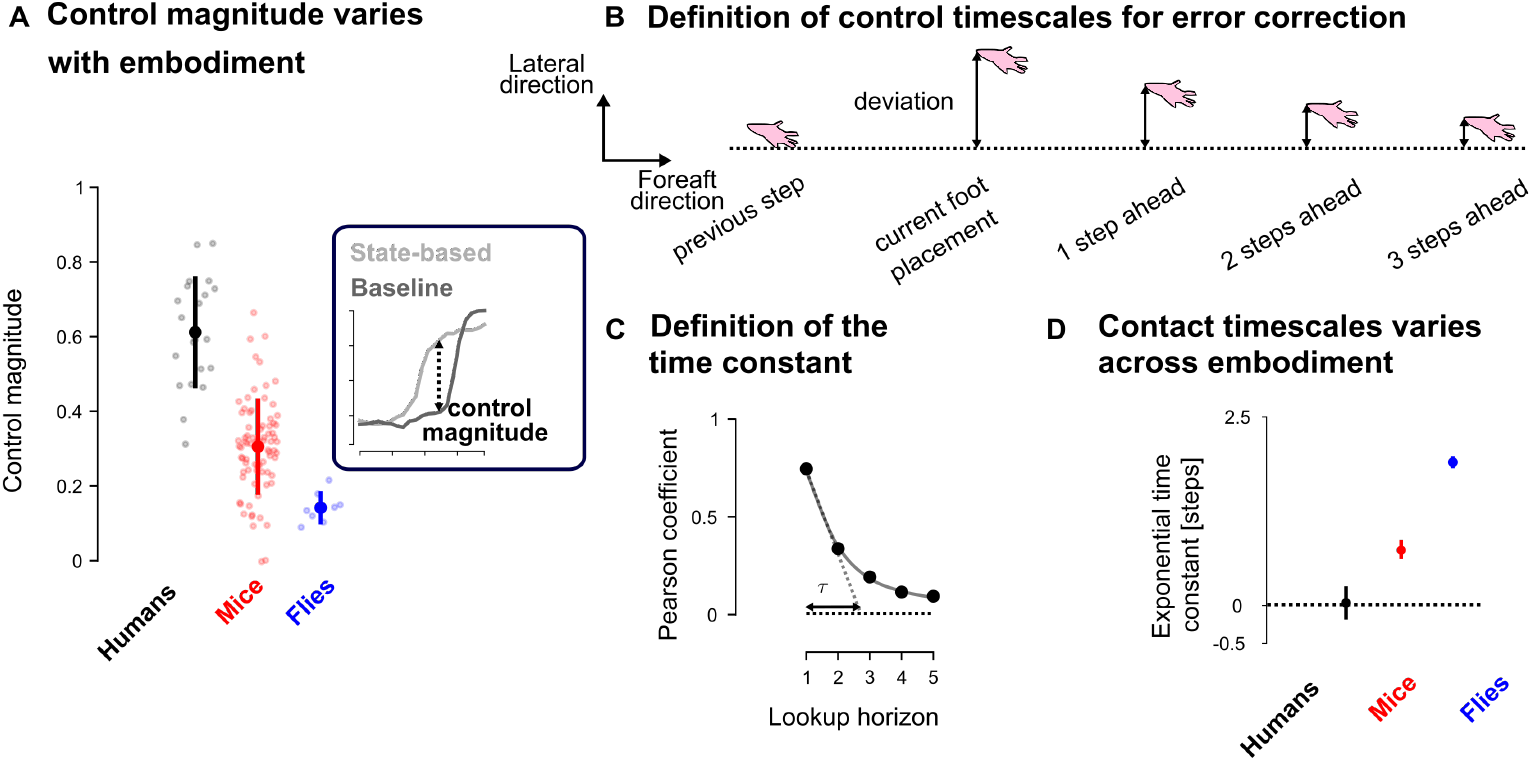
Magnitude and timescale of the control signatures vary across embodiments. **A** Group median and interquartile range of the control magnitude for the lateral foot placements of humans legs (black), mice front legs (red) and flies front legs (blue) at the initiation of the swing phase of the contact foot. The control magnitude is defined as the difference between the amount of variance explained by the body state-based and the baseline prediction (see insert). **B** Schematic representation of the variations in the lateral foot placement as a function of time. We computed the correlation between the current lateral foot placement and the subsequent lateral foot placements. The Pearson coefficient of correlations exponentially decayed with the look up horizon as schematized in panel **C**. We characterized this exponential decay by the exponential time constant, *τ*. **D** Exponential time constants in humans, mice and flies (maximum across legs for each species). The error bar represent the centered 90% confidence interval based on bootstrap resampling.

While we found that the body state error-dependent foot placement control magnitude was lower for more passively stable animals, this did not tell us about the *control timescale* i.e. over how many foot contacts the error was corrected. To capture that control timescale, we computed the correlations between a given lateral foot placement of a limb and its subsequent lateral foot placements (Figure 4B). These correlations, captured by the Pearson coefficient, exponentially decayed with time (Figure 4C, illustrative data). We extracted the maximal exponential time constant (*τ* in Figure 4C), across limbs, for each species and observed a gradient across them (Figure 4D) with the smallest time constants observed in humans (0.035±0.43), followed by mice (0.73±0.2) and flies (1.89±0.11). These results suggested that the timescale over which errors are corrected vary with many-legged embodiment – the correction timescale increased with the number of legs.

Thus, we found that the magnitude and timescale of the lateral foot placement control signatures vary with many-legged embodiment; with larger number of limbs, the magnitude decreased, and the timescale became increasingly multi-step.

### Signatures of direction- and leg-specific modular control of foot placement

We have discovered shared signatures of body state error-dependent foot placement control from locomotor variability in many-legged animals, and quantified how the control magnitude and timescale vary with embodiment. With the availability of multiple legs to execute the same functional goal, namely stable locomotion in the presence of sensorimotor noise, animals may use a more or less centralized control strategy i.e. they may use different legs or muscle groups for executing modular control functions [66]. Here, we analyzed the locomotor variability to test the existence of such leg-specific or direction-specific modular control.

We hypothesized that, with the availability of multiple legs for control, medial and lateral body state errors relative to the stance foot i.e. errors that are directed towards and away from the stance foot respectively, would be controlled differently (see Figure 5A). The rationale for this hypothesis is that medial errors in the body state near the frontmost limbs directed towards the stance foot can be corrected by the impedance of the stance foot itself, unlike lateral errors directed away from the stance foot and towards the swinging foot. To test this hypothesis, we computed the feedback gains associated with medial and lateral body errors separately (see *Methods*) and compared them at the beginning of the swing phase of the foot being placed. Medial and lateral feedback gains were similar in humans (Figure 5B left, Wilcoxon signed-rank, *p* = 0.13, *d* = 0.38) but differed in mice (Figure 5B center, Wilcoxon signed-rank, *p <* 0.05, *d* = 1.57) and flies (Figure 5B right, Wilcoxon signed-rank, *p <* 0.05, *d* = 1.20), where the corrections were larger for lateral than for medial errors.

**Figure 5:**
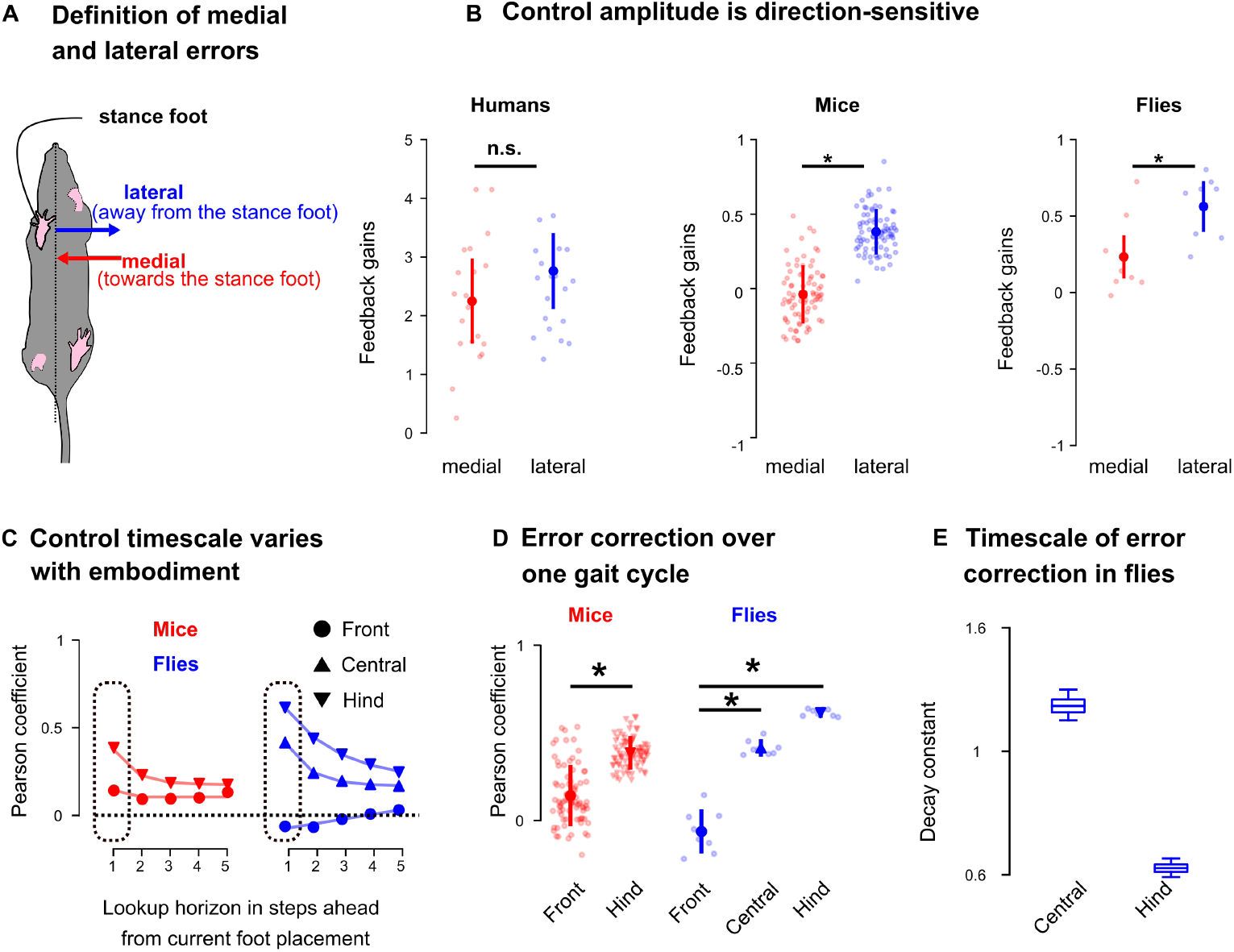
Foot placement control signatures reveal direction-specific and limb-specific modularity across species. **A** We illustrate the definition of lateral (blue) and medial (red) errors respectively as the errors directed away and towards the stance foot. We independently fitted models for lateral and medial errors and compared their foot placement control gains. **B** Group median and interquartile range of the lateral (blue) and medial (red) foot placement control gains for humans, mice and flies at the beginning of the swing phase of the contact foot. **C** Pearson correlation coefficients as a function of lookup horizon for the different limbs in mice (left, red) and flies (right, blue), group median. **D** Group median and interquartile range of the initial Pearson correlation coefficient (see dashed rectangle in panel C) for the different pairs of limbs in mice and flies. **E** Timescale of the error corrections for central and hind limb in flies, the error bars capture the 90% confidence interval. ∗ : *p <* 0.05

Within a given embodiment, we posit that different pairs of limbs could have different control strategies; the rationale for this hypothesis is that in quadrupedal animals the front and hind limbs are innervated by different spinal circuits and therefore may exhibit distinct behavioral control signatures [17]. To test the existence of such leg-specific differences in control strategies, we compared the mapping between the body state errors and lateral foot placement across different limb pairs – from anterior to posterior as defined by the movement direction – over multiple gait cycles (Figure 5C). We observed that the correlations over a single gait cycle (see dashed inserts in Figure 5C) varied across legs; larger values were observed for more posterior limbs in both mice (Figure 5D red, Wilcoxon signed-rank test, *p <* 0.05, *d* = 1.37) and flies (Figure 5D blue, Kruskal-Wallis with posthoc Dunn’s test and FDR (false detection rate) corrections, front-central : *p <* 0.05, *d* = 1.88, front-hind : *p <* 0.05, *d* = 1.94). The exponential time constant associated with the decay of these correlations also varied across limbs in flies where the hind limbs had a smaller constant, i.e. a slower correction timescale, than the central limbs (Figure 5E). These results suggest that the control strategies for lateral foot placement in different limbs of flies and mice operate with different timescales – the more posterior pairs of limbs operate at slower timescales and have persistent control across multiple contacts.

Thus, we found that the control strategies used for foot placement by many-legged species reveal direction- and leg-specific modular control, exhibiting variation with error direction along the medial-lateral axis, and with limb pairs along the anterior-posterior axis relative to movement direction. We find that both the control gains and timescales exhibit such leg- and direction-specific modularity.

## Discussion

We put forth a unified feedforward-feedback control structure underlying foot placement for stable locomotion in any legged animal. Based on this control structure, we develop a data-driven model to discover signatures of control from natural locomotor variability across species. Using this model, we discover signatures of body state error-dependent locomotor control in flies and mice, previously only shown to exist in humans. We find that the urgency of foot placement control, reflected in the control magnitude and timescale, decreased in more passively stable embodiment. Foot placement control is executed modularly in many-legged animals as illustrated by the use of limb- and direction-specific gains and timescales. These findings have implications for understanding the neuromechanics of animal locomotion, uncovering the neural basis of stabilizing control, developing cross-species phenotypes for locomotor control, and bio-inspired legged robots.

### Foot placement control signatures in flies, mice, and humans

We discovered signatures of body state error-dependent control from unperturbed locomotor variability in multi-legged animals, a phenomenon previously only observed in humans [64, 3, 4]. This is the first cross-species characterization of body state error-dependent foot placement control, and importantly, it does so by unifying velocity-dependent and body state error-dependent contributions to control. Our feedforward control module captures the stereotypical gait patterns of different species and how they co-vary with movement velocity. Our feedback control module, which captures the body-state error-dependent corrective foot placements, echoes previous work highlighting the importance of lateral foot placement for stable locomotion in bipeds [18, 3], and provides the first cross-species body state error-dependent signatures of foot placement control, previously posited in theoretical models of legged locomotion [36, 37]. Moreover, while previous work reported similarities in the structure of kinematic variability between mice and flies [67, 68], our work demonstrates how this variability relates to stability by correlating foot placement to recent body state errors. Here, by eliminating the velocity-dependent component of foot placement, our data-driven model is able to avoid erroneous interpretation of body state error-dependent foot placement in the fore-aft direction (Figure 3C), supported by theoretical modeling [59]. Overall, our findings are consistent with body state error-dependent lateral foot placement control across species.

### New empirical support for the control structure underlying foot placement

Using a data-driven model inferred from natural locomotor variability, we evaluated two competing hypothesized control structures for legged locomotion. The first structure posits purely velocity-dependent feedforward control (Figure 1A, red), while the second (ours) also posits body-state error-dependent feedback control in the vicinity of the feedforward controller (Figure 1A, red+blue), contrasting previous data-driven studies that defined such feedback control in the vicinity of the average behavior [3, 58, 59]. Our data-driven signatures in humans, mice, and flies are consistent with the feedforward-feedback control structure. Although both theorized control structures explain the data-driven velocity-dependent component of foot placement location and timing, only the feedforward-feedback model can explain the lateral foot placement variability (Figure 3B). Thus, we provide cross-species empirical support consistent with the hypothesis that velocity-dependent control is insufficient for lateral stability [36, 37]. In this work, we more clearly link the body state error-dependent feedback control signatures to their stabilizing effect on the body by identifying a positive correlation between corrective foot placements on one step and subsequent error reduction for the body state on the next step. While our data-driven behavioral signatures alone do not guarantee the existence of causal body state error-dependent control during locomotion, when viewed in combination with previous theoretical work [36, 37, 18, 38], it puts forth a more compelling case by providing missing empirical support.

### Neuromechanical basis of foot placement control

The signatures of foot placement control we discovered could help investigate the neuromechanical basis of stable locomotion across species. The hypothesized feedforward-feedback control structure suggests distinct, yet interacting mechanisms for velocity-dependent and body state error-dependent control. The signatures of velocity-dependent fore-aft foot placement and contact timing (Figure 2) aligns with the periodic velocity-dependent coordination afforded by pattern-generating neural circuits [6, 17, 46]. The lateral body state error-dependent foot placement (Figure 3) suggests an additional feedback mechanism, which we speculate could map to sensory feedback-guided steering. Such steering has been found in mice [50], flies [69, 70], and humans [71, 72]. We speculate that these two mechanisms interact based on our feedforward-feedback control structure, because the feedback module relies on information about the nominal body state from the feedforward module (Figure 3B). We speculate that such body state error estimation in mice and humans could rely on vestibular information [50, 51, 49] mediated through supraspinal neural centers [73, 74, 75, 69, 76, 77, 78], whose disruption affects locomotor control [48, 79]. The observed cross-species differences in foot placement control amplitudes and timescales (Figure 4) suggest links to the differences in neuromechanical embodiment. While our data-driven models alone cannot definitively distinguish between the contributions of biomechanics, neural circuitry, or both, to the observed differences in foot placement control across species, these findings open exciting avenues for future research. For example, combining our approach with mechanical [45, 80] or neural perturbations [50] could help isolate the neural versus biomechanical contributions to control. Alternatively, our data-driven model could be combined with physics-based imitation learning to posit the neuromechanical basis of the observed signatures in simulation [81, 82]. These findings can also inform the development of cross-embodiment control strategies in legged robots [83], by providing bio-inspiration for the selection of neuromechanical control primitives (Figure 5). Overall, understanding how foot placement control strategies change with neuromechanical embodiment across species for a shared functional goal can provide insight into the co-evolution of biomechanics and neural control.

### Inferring control from intrinsic versus external perturbations

Here, we investigate the control strategies hidden in natural locomotor variability that ensure stability in the presence of intrinsic sensorimotor noise-like perturbations as opposed to externally applied perturbations. Such a “default controller” [84] to handle perturbations due to intrinsic errors is ethological, because all animals likely have a degree of unavoidable sensorimotor noise that continuously perturbs their movements [85, 86], which they must remain stable to. Indeed, in humans, studying this default controller hidden in natural variability has been important for understanding foot placement control [84, 5, 3], revealing that this controller is robust enough to handle larger perturbations [5, 58]. Inferring control from natural variability also offers a link to numerous mathematical formulations of stability [87], following the example of how these formulations have been linked to variability in human locomotion [4]. For instance, the signatures of body state error-dependent feedback control in multi-legged animals, as discovered here, is reminiscent of the center of mass-based control in humans and suggests that stability formulations developed for abstract inverted pendulum models [18] or more detailed robotic models [88] could be applied across species. Furthermore, our findings of control signature timescales in more inherently stable animals aligns with previous work on the impact of leg number and configuration on dynamic stability [37]. Finally, our findings of embodiment-dependent control timescales can be linked to theoretical studies that posit the number of steps required to reach a stable body state [89, 65, 90, 91] which is related to falls in humans [92]. By deriving control signatures from natural variability, we can bridge the gap between theoretical models and natural locomotion across species.

### Weaknesses and strengths of studying control from natural variability

Our work provides a way to compare control strategies across species for a shared ethological goal: foot placement control in the presence of intrinsic sensorimotor noise. Investigating animal locomotion in such an ethological setting has both limitations and strengths. One key limitation is the inability to know, from behavior alone, the precise nature of the noise-like input which results in errors to be controlled. Here, we hypothesized that the observed foot placement variability is the result of feedforward-feedback control in response to intrinsic sensorimotor noise (Figure 1A), which has some theoretical basis [86, 5], but confirming whether this noise is motor, sensory, or neural in origin is challenging with theory and behavior alone [93]. Similarly we hypothesized that the differences we observed across species (Figure 4) are related to the differences in their neuromechanical embodiment, but our natural variability-based data-driven models cannot discriminate between biomechanical and neural contributions to these differences. These challenges are an opportunity for future neuromechanics research to uncover the source of intrinsic sensorimotor errors and that of the cross-species differences in foot placement control. Even if we only analyzed walking gaits, i.e. gaits for which there is no flight phase, we suspect that our approach could generalize to faster gaits in mice and flies, as it did in humans [5]. However, the reliance on body state error-based control could significantly decrease at high speeds [94, 95, 96] because of the lack of time to process delayed sensory inputs. There are also strengths to studying the control strategies underlying natural variability, such as the discovery of ethological control principles, the relevance to easily quantifiable clinical biomarkers, and the ease of reproducibility. Since all animals have some degree of intrinsic sensorimotor noise, inferring control strategies from natural variability is ethological and representative of the animal’s natural control repertoire [97, 98]. Population differences in human locomotor variability are used as biomarkers for neuromotor disorders that impair balance, such as Parkinson’s disease [60]. Here, we have shown that neuroscientifically accessible animal models like flies and mice exhibit human-like signatures of control hidden in variability, thus encouraging the study of these control signatures as potential cross-species biomarkers. Our approach is easily reproducible as it is applicable to any multi-legged animal (see *Methods*), builds on open source video processing pipelines [99, 100], and does not require specialized equipment other than a camera placed such that foot contacts are discernible. Moreover, as locomotion is a behavior that is readily exhibited by animals, our approach can enable post-hoc analyses of large-scale animal behavior datasets [48].

## Conclusion

This work establishes three key findings: (i) cross-species signatures of body state error-dependent foot placement control across species, (ii) how these control signatures change with neuromechanical embodiment, and (iii) data-driven signatures consistent with a combination of velocity-dependent and body state error-dependent control, and inconsistent with purely velocity-dependent control in flies, mice, and humans. These findings provide a starting point for future investigations into the neuromechanical basis of foot placement control across species.

## Methods

### Mathematical formulation of feedforward-feedback foot placement control

We hypothesize that foot placement control in many-legged animals is generated by a feedforward-feedback control structure (Figure 1A) composed of a velocity-dependent feedforward module and a body state error-dependent feedback module. This control structure outputs the net observed foot placement location and timing as the resultant of the contribution of its two control modules (Figure 1D): the feedforward module contributes the foot placement prescribed by the canonical gait patterns [67] while the feedback module contributes body state error-based corrective adjustments of foot placement [3]. These modules operate at different timescales and are organized in a *hierarchical* manner such that each of them has access to some privileged information which is not available to the other. The input to the feedforward control module is the average velocity during the last gait cycle 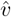 while the inputs to the feedback control module are the body state deviations from the stereotypical species-specific gait pattern *Q*^∗^, expressed as phase-dependent body state errors Δ*Q*(*ϕ*) = *Q*(*ϕ*) − *Q*^∗^(*ϕ*), where *ϕ* is the relative gait fraction defined as the ratio between the duration since the beginning of the last gait cycle and the total duration of that gait cycle. For each step, the net observed foot placement was expressed as the sum of the individual contributions from the two control modules

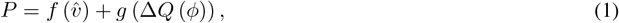

where functions *f* and *g* are respectively the contributions of the feedforward and feedback control modules. We used the data from naturalistic locomotion of different legged species to investigate signatures of the hypothesized control structure across species.

### Inference of control signatures from locomotion data

We combined the hypothesized control structure underlying legged locomotion with a data-driven model to investigate signatures of feedforward and feedback foot placement control. First, we investigated the velocity-dependent component of foot placement in the fore-aft and lateral directions. To do this, we generalized the methodology used in humans to many-legged embodiment [63]: for each gait cycle, we defined the movement velocity as the average fore-aft velocity of the body during the last gait cycle. We observed that the relationships between the movement velocity and the fore-aft and lateral foot placement were linear (Figure 2A-C) and characterized this relationship with the model

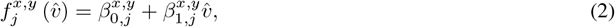

where 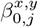 and 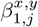 are scalar model parameters capturing the relationships for the *jth* leg along the fore-aft (*x*) and lateral (*y*) direction. We used individual models for each animal subject and for each limb to capture how that limb’s step length and width were modulated by the average velocity. We observed that the relationships between movement velocity and the contact timing followed an exponential function (Figure 2E-G) and characterized this relationship with the model

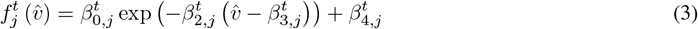

where 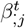 are scalar model parameters capturing the relationship for the *jth* leg.

To investigate the behavioral control signatures of the hypothesized feedback module, we defined the body state errors as the difference between the body state kinematics (positions and velocities) during the last gait cycle *Q* (*ϕ*) and the velocity-dependent body states as specified by the stereotypical gait pattern 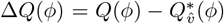, where 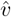 is the average velocity over that gait cycle. We defined the corrective foot placements as the difference between the net observed foot placement *P* and the ones predicted by the feedforward control module 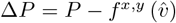. We observed linear relationships between the body state errors and the corrective foot placements (Figure 3B), which is theoretically valid for small errors, because continuous and differentiable nonlinear functions can be approximated as linear for infinitesimal changes. We investigated these relationships for the *jth* leg by the model

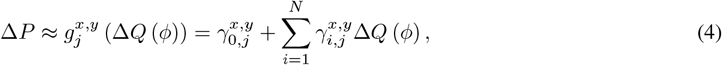

where 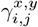 are scalar model parameters and *N* is the dimensionality of deviation vector.

We investigated the behavioral signatures of the feedback control module across species by comparing their amplitude and timescale. The control amplitudes were quantified as the maximal individual differences between the amount of variance explained by the body state error-dependent feedback model (blue traces in Figure 3B) and that of a baseline model predicting the variability in foot placement from the foot kinematics itself (red traces in Figure 3B). This baseline model, expressed as a function of the deviations in body kinematics 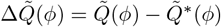, was formalized as

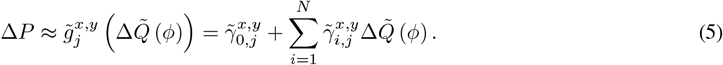

We then investigated the control timescales by correlating successive lateral positions of each foot over multiples gait cycles. The limb-specificity of the feedback control module was investigated by separating the medial and lateral errors and using the feedback model described above. Consistent with anatomical definitions, these direction-dependent errors 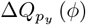 are defined as follows: medial errors are directed towards the body while lateral are directed away from the body (Figure 5A.)

We assessed the locomotion stability using Poincare return maps applied to the deviations in body kinematics. We defined these return maps based on the following relationship

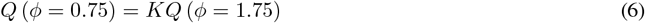

where K is a scalar matrix evaluated based on the data. The eigenvalues of this matrix, *λ*_*i*_ were computed for each species and the spectral radius defined as *ρ* (*K*) = max_*i*_ |*λ*_*i*_| was reported to characterize the stability of the system.

We computed the variations in body state errors around contact as the difference between the averaged body errors during the half-gait cycle before and after contact based on the following relationship

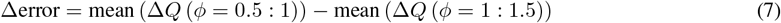

### Datasets used to investigate foot placement control across species

The contributions of the feedforward and feedback control modules can be inferred for any dataset for which kinematics data of the body and feet are available. To demonstrate this flexibility, we chose to use existing open source datasets of flies, mice and humans during natural locomotion collected across different research groups.

The fly locomotion dataset was collected by DeAngelis and colleagues [68] with a Point Grey camera (Flea3-U3-13Y3M-C, 150Hz) when 8 groups of 12 to 15 flies (*Drosophila melanogaster*) were evolving in an open-field arena during 1.1h. The data was segmented into individual locomotion bouts and the body and feet positions, as well as the individual foot contacts were extracted by the original authors using a custom-built markerless motion capture method (see [68] for more details). We used the feet and the head markers in our analyses and only considered the segments during which the animals were not stopping. The mouse locomotion dataset was collected by Klibaite and colleagues with a Point Grey greyscale camera (12-bit greyscale, 1280 × 1024 pixels, 80Hz) when mice (C57BL/6J) were moving in an open-field arena on 4 consecutive days, 20 minutes each day. The positions of the different body parts were extracted using LEAP [100], an open source markerless pose estimation software. We used the feet, nose, and base of the tail markers for our analyses and only considered the data segments labeled as locomotion. The human locomotion dataset was collected by Camargo and colleagues [101] using 32 motion capture markers placed according to the Helen Hayes Hospital marker set (Vicon. Ltd, Oxford, UK). The data were collected at a frequency of 200Hz and was filtered using a 4th order zero-lag Butterworth filter (cutoff frequency 6Hz) during overground walking. We used the pelvis position, defined as the average position of the four hip markers, and the foot position, approximated by the position of the heel markers. We also used another human locomotion dataset collected by Wang and Srinivasan [3] for some analyzes, the data collection procedure was the same as that of the Camargo dataset and we used the same set of markers.

### Processing pipeline for obtaining the inputs and outputs of the foot placement controller

In this section, we describe how we processed the raw data described above to transform them into the inputs and outputs to the feedforward and feedback control modules. We sought to use an identical processing pipeline for all the datasets. In line with this, the steps outlined below were applied to all the datasets and species, unless otherwise stated.

We were interested in studying the straight-line locomotion bouts since foot placement control has only been characterized during straight-line locomotion in previous theoretical and human literature [2, 18, 3]. We therefore selected the locomotion bouts for which the body orientation defined as the direction of the instantaneous velocity vector in the horizontal plane did not vary by more than 30 degrees. We then rotated these straight-line locomotion bouts to align the heading direction across bouts, since the raw data contained locomotion bouts that can have different orientations in the world coordinates. We next identified the timing of each foot contact to extract the contact locations and segmented the gait cycles. For the datasets in which the contact timings were not provided, we computed the distance between the feet and body markers in the horizontal plane and defined the beginning and end of the stance phases as the maxima and minima of that distance [102]. The prediction of contact locations obtained with this method were similar to those obtained with a velocity-thresholding of the foot marker (Figures S1 and S2), while being less sensitive to noise. We defined the contact locations, *P* = [*p*_*x*_, *p*_*y*_]^*T*^ as the average positions of the foot markers during their stance phases and the contact timings as the time at which the swing to stance transition occurs. We used the contact initiations i.e. the beginning of the stance phases to perform the gait segmentation, defining a gait cycle as the interval between two successive contact initiations of a same leg. We performed this gait segmentation independently for each limb and grouped the gait cycles associated with the same limb. We compared the results obtained with a gait segmentation based on successive contact terminations of a same leg and did not observe any difference (Figure S3).

Because the inputs to the feedback control module are posited to be phase-dependent Δ*Q* (*ϕ*), we temporally normalized the segmented gait cycles by linearly interpolated their marker positions and velocities onto a fixed set of discrete phase *ϕ*. The number of phases for each species was set to the average number of frames per gait cycle in the raw data. We validated this approach by comparing the predicted phase to the Phaser algorithm [103] which estimates the gait phase by combining the individual contributions of each leg; there was no systematic bias or difference between the predictions of these two approaches (Figure S4). Therefore, body and foot states are represented relative to phase *ϕ*, with *ϕ* = 0 corresponding to the beginning of the gait cycle and *ϕ* = 1 to the end of the gait cycle, which is when the contact corresponding to foot placement occurs. We then spatially normalized the marker positions and the contact locations by subtracting the last contact location of the front limb belonging to the opposite group of limbs. We defined the opposite group of limbs based on the canonical gait patterns whereby two groups of limbs are moving in opposite phase during walking (filled and empty rectangles in Figure 1B).

### Statistical analyses

All the statistics were performed using Python3 (*numpy, scipy*, and *scikit_posthoc*). We used non-parametric tests whenever the normality condition, assessed with the Shapiro-Wilk test, was not met for at least one of the species. We used the Kruskal-Wallis followed by Dunn’s posthoc with FDR (false detection rate) corrections for multiple comparisons. We calculated and reported the effect size, quantified by the Cohen’s d for each comparison of two variables. We reported significance when the p value felt below 0.05.

We used the chatterjee-xi coefficient [104] to investigate correlations between pairs of variables. Chatterjee’s coefficient of correlation was chosen over Pearson’s coefficient as it is not restricted to monotonic trends. The chatterjee-xi test from *scipy* was used to test whether the variables were independent. We studied the specific impact of movement velocity on the step length and width (Figure 2) by fitting individual linear regressions, motivated by previous studies [63, 10]. We used the function *scipy*.*stats*.*linregress* in Python to fit these regressions and kept track of the individual slopes and p values associated with each of these.

To infer the feedback control module (Figure 3), we obtained linear models between the body errors Δ*Q* (*ϕ*) and the foot placement deviations Δ*P*. The best fit model parameters, obtained by the minimization of the mean squared errors, were given by 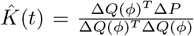 and the coefficient of determinations were given by 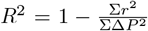, where 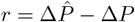 are the residual errors associated with the multiple linear regression.

To investigate whether errors were corrected over multiple gait cycles (Figure 4), we investigated the correlations between successive lateral foot placements for each leg. We computed these correlations between lateral foot placements at time *t* and those at time *t* + *N* where *N* is an integer value capturing the lookup horizon. We then obtained the exponential decay capturing the variation of the Pearson coefficient with the lookup horizon. We used bootstrap resampling (n=10 000) to evaluate the parameters of this exponential decay and reported the 90% confidence interval for each of the parameters we studied.

## Acknowledgments

The authors thank Manoj Srinivasan, Hari Teja Kalidindi, and the members of the Seethapathi Motor Control group for their feedback on an earlier version of this manuscript. Antoine De Comite is supported by a postdoctoral fellowship of the K. Lisa Yang Integrative Computational Neuroscience (ICoN) Center at MIT.

## S1 Methodological details for calculating gait and phase events

### Velocity-based contact detection method

In the main paper, we detected the timing of contact initiation and termination by using the maxima and minima of the fore-aft distance between the feet and the body markers. This method is prone to systematic bias in the estimation of contact timing if there is leg retraction prior to contact initiation. Leg retraction consists of a rearward movement of the leg just before heel strike. In the presence of such leg retraction, our method will detect a maximum when leg retraction starts, which does not necessarily correspond to contact initiation. To validate our contact detection method and guarantee that leg retraction was not systematically biasing our contact detection, we compared our method’s prediction to that of another method which also considers the foot velocity for contact detection, seeking to avoid artifacts linked to leg retraction [105]. Briefly, this second method uses a threshold on velocity alongside the maximum distance in the fore-aft direction to identify contact initiation. For each detected maximum of the fore-aft distance (i.e. putative foot strike), we search the first time at which the foot velocity drops below a threshold. We observed that the fore-aft and lateral location of individual foot contact detected with both methods, defined as the average position of the foot during the stance phase, are extremely similar (Figure S1) and that the predictions of the body state error-based feedback control module are unaltered when the velocity-based method is used (Figure S2). However, the velocity-based method introduces some noise in the estimation of contact location (as illustrated in Figure S1A), we therefore used the position method in the main manuscript.

**Figure S1:**
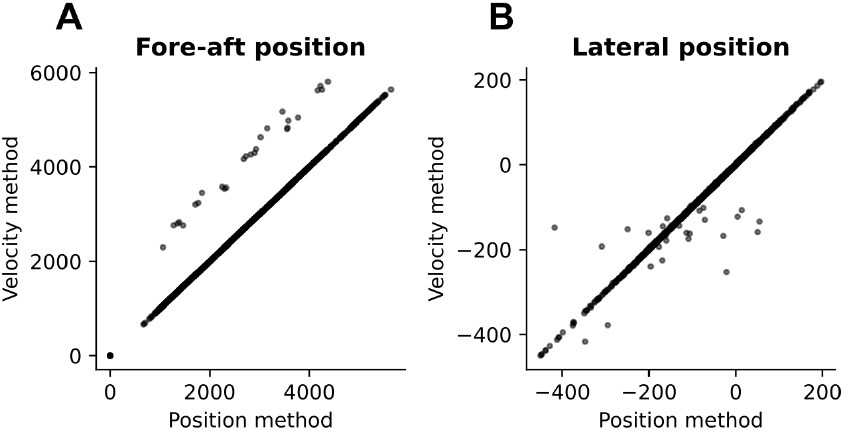
Fore-aft (**A**) and lateral (**B**) contact location predictions for the position-only method (horizontal axis) and the velocity-based method (vertical axis) in humans.

**Figure S2:**
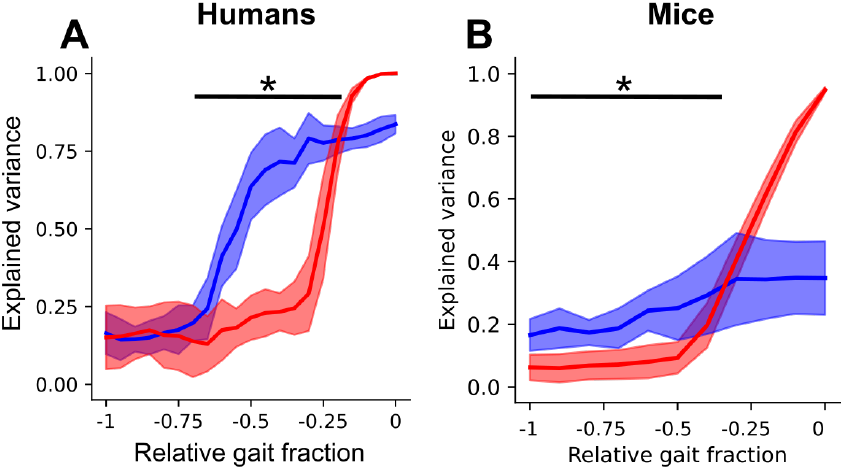
Explained variance (R^2^, median and interquartile range) obtained for the body state-based (blue) and baseline (red) predictions of the lateral deviations in foot placement for humans (**A**) and mice (**B**) using the velocity-based contact detection method.

### Gait segmentation

When investigating the signatures of body state error-dependent foot placement, we used a gait segmentation based on the initiation of foot contact, i.e. touchdown-to-touchdown gait segmentation. To investigate whether this choice of gait segmentation altered our interpretation of the signatures, we conducted the same analyses by using a gait segmentation based on the initiation of the swing phase, i.e. liftoff-to-liftoff gait segmentation. This segmentation based on liftoff reveals qualitatively similar signatures as reported in the main manuscript (Figure S3). We therefore did not observe any effect of the choice of gait events used for segmentation on the finding of the body state error-dependent foot placement control signatures.

**Figure S3:**
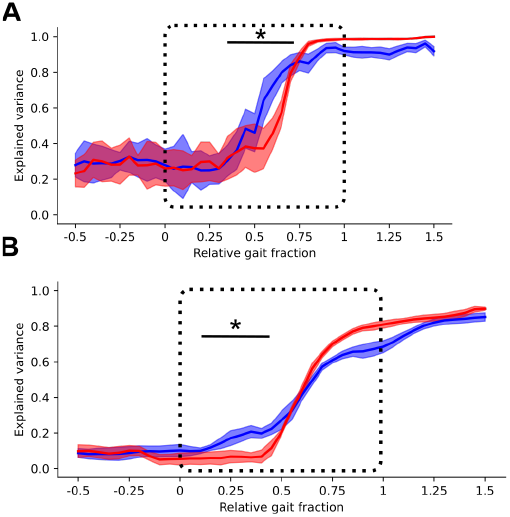
Explained variance (R^2^, median and interquartile range) obtained for the body state-based (blue) and baseline (red) predictions of lateral deviations in foot placement in humans (**A**) and flies (**B**) for a liftoff-to-liftoff gait segmentation. The black dashed box represents the phase window shown in Figure 3 of the main manuscript. The stars indicate the phase domain where the body state-based model outperformed the baseline (2 samples Kolmogorov-Smirnov tests).

### Phase estimation

We compared the method we used to estimate the relative gait fraction to a another method that estimates the locomotion phase with a dedicated algorithm [103]. Briefly, this method first computes the Hilbert transform of the fore-aft distance of each individual foot. Then, extracts the *protophase* of each leg as the complex argument of this Hilbert transform. Finally, the individual phases are combined in a single phase estimate using dimensionality reduction techniques. We compared the phase estimations obtained with this method (*oscillators-based method*) to that obtained by linearly interpolating the time vector between two successive contacts of the same leg (*time-based method*) and did not observe any systematic differences across species (Figure S4A-C).

**Figure S4:**
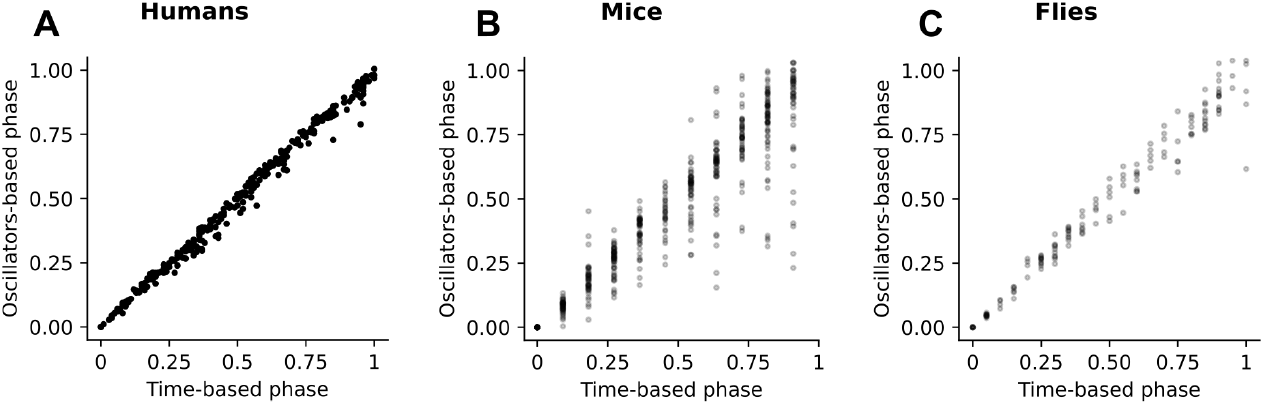
Comparison of the relative gait fraction obtained by the time-based method (horizontal axis) and the oscillators-based method (vertical axis) for humans (**A**), mice (**B**), and flies (**C**).

## S2 Analyzing the relationship between foot placement and future error reduction

We analyzed the potential stabilizing effects of the foot placements by correlating them to the error reduction on the next step. We defined the error reduction as the difference between the average body state errors during the half gait cycle following contact and the average body state error during the half gait cycle preceding contact. In the main manuscript, we reported the statistics associated with the linear correlations between lateral foot placement and lateral error reduction (Figure S5D-F). The fore-aft deviations in foot placement, on the other hand, were not correlated with fore-aft error reduction (Figure S5A-C). This finding is consistent with our conclusion that there is body state-dependent error correction in the lateral but not in the fore-aft direction.

**Figure S5:**
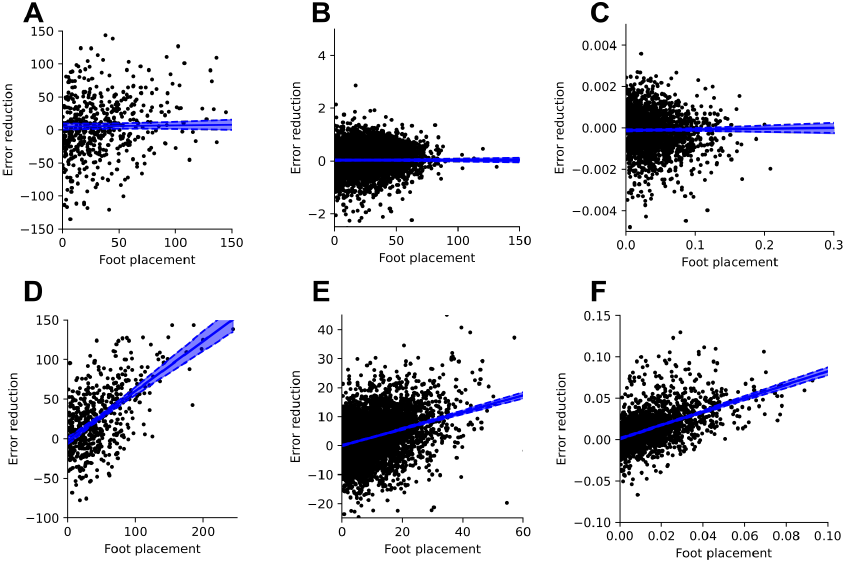
Error reduction as a function of foot placement deviation for the fore-aft (top row) and lateral (bottom row) directions for humans (leftmost column), mice (central column), and flies (rightmost column). The associated linear regression and their corresponding 95% confidence intervals are represented in blue full lines and shaded areas, respectively.

## S3 Quasi-experimental causal inference of foot placement control signatures

Scientific domains like economics [106], neuroscience and behavior [107] use quasi-experimental methods for causal inference using observational data. Inspired by the *differences-in-differences* method, we quantify the effect of body state-dependent foot placement control during the previous step on the body state error reduction on the next step. For a given small range of positive body-state errors on the previous step, we defined the treatment and the no-treatment groups as the trials with positive and negative body state-dependent foot placement errors respectively. We then compared the error reduction associated with these two groups and reported those for which the distribution of error reduction is significantly greater than zero, corresponding to a reduction in body state-error associated with that specific contact. We found a significant effect in the treatment group for all three species (Figure S6A-C).

The main challenge in extending these quasi-experimental approaches to a new domain such as ours is determining what the treatment variable is. In the case presented here, we considered the presence of body state-dependent deviations in foot placement as treatment and performed the subsequent analyses accordingly. In future extensions of this work that include mechanical or neural perturbations, these interventions could be considered treatment variables, enabling a quantitative test of the putative causal relationship between body state errors and foot placement corrections.

**Figure S6:**
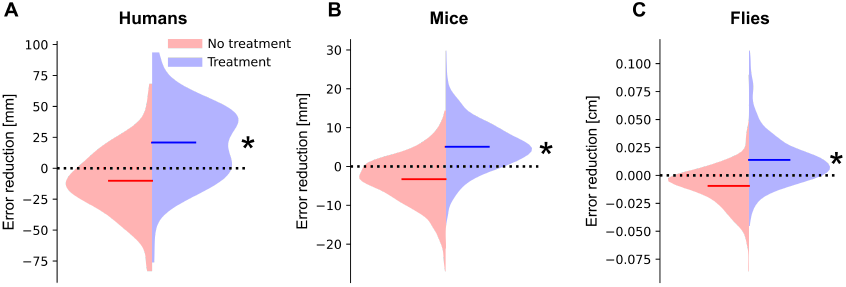
Distributions of error-reduction for the treatment (blue) and no-treatment (red) groups for humans **A**, mice **B**, and flies **C**. The dashed black horizontal lines represent an absence of error reduction, positive and negative values respectively capture reduction and increase in body state errors. Significance is reported for the distributions statistically larger than zero. *: *p <* 0.05

## Notes

### Competing Interest Statement

The authors have declared no competing interest.

### Summary of Updates

Now includes supplementary materials for detailed methods description

## References

[1] MH Raibert, HB Brown, M Chepponis, and E Hastings. Dynamically stable legged locomotion research: December 1, 1980-september 30, 1981. Technical report, Carnegie Mellon University, 1981.

[2] MA Townsend. Biped gait stabilization via foot placement. Journal of Biomechanics, 18(1):21–38, 1985.

[3] Y Wang and M Srinivasan. Stepping in the direction of the fall: the next foot placement can be predicted from current upper body state in steady-state walking. Biology Letters, 10, 2014.

[4] SM Bruijn and JH van Dieën. Control of human gait stability through foot placement. Journal of the Royal Society Interface, 15, 2018.

[5] N Seethapathi and M Srinivasan. Step-to-step variations in human running reveal how humans run without falling. eLife, page e38371, 2019.

[6] AJ Ijspeert. Central pattern generators for locomotion control in animals and robots: A review. Neural networks, 21:642–653, 2008.

[7] S Grillner and A El Manira. Current principles of motor control, with a special reference to vertebrate locomotion. Physiological Reviews, 100:271–320, 2019.

[8] P Cisek and BY Hayden. Neuroscience needs evolution. Philosophical Transactions of the Royal Society B, 377:20200518, 2001.

[9] JW Krakauer, AA Ghazanfar, A Gomez-Martin, MA MacIver, and D Poeppel. Neuroscience needs behavior: Correcting a reductionist bias. Neuron, 93:480–490, 2017.

[10] NC Heglund, CR Taylor, and TA McMahon. Scaling stride frequency and gait to animal size : mice to horses. Science, 186:1112–1113, 1974.

[11] RM Alexander and AS Jayes. A dynamic similarity hypothesis for the gaits of quadrupedal mammals. Journal of Zoology, 201:135–152, 1983.

[12] C Bellardita and O Kiehn. Phenotypic characterization of speed-associated gait changes in mice reveals modular organization of locomotor networks. Current Biology, 25:1426–1436, 2015.

[13] C Chun, T Biswas, and V Bhandawat. Drosophila uses a tripod gait across all walking speed, and the geometry of the tripod is important for speed control. eLife, page e65878, 2021.

[14] JE Bertram and A Ruina. Multiple walking speed-frequency relations are predicted by constrained optimization. Journal of theoretical Biology, 209:445–453, 2001.

[15] JA Nirodi. Universal features in panarthropod inter-limb coordination during forward walking. Integrative and Comparative Biology, 61:710–722, 2021.

[16] C Mantziaris, T Bockemuhl, and A Buschges. Central pattern generating networks in insect locomotion. Developmental Neurobiology, 80:16–30, 2020.

[17] F Delcomyn. Neural basis of rhythmic behavior in animals. Science, 210:492–498, 1980.

[18] AL Hof, MG. J Gazendam, and WE Sinke. The condition for dynamic stability. Journal of Biomechanics, 38(1):1–8, 2005.

[19] G Courtine, AM De Nunzio, M Schmid, Berettan MV, and M Schieppati. Stance- and locomotion-dependent processing of vibration-induced proprioceptive inflow from multiple muscles in humans. Journal of Neurophysiology, 97:772–779, 2007.

[20] DC Roden-Reynolds, MH Walker, CR Wasserman, and JC Dean. Hip proprioceptive feedback influences the control of mediolateral stability during human walking. Journal of Neurophysiology, 114:2220–2229, 2015.

[21] LR Bent, TI Inglis, and BJ McFadyen. When is vestibular information important during walking ? Journal of Neurophysiology, 92:1269–1275, 2004.

[22] RC Fitzpatrick, DL Wardman, and JL Taylor. Effects of galvanic vestibular stimulation during human walking. Journal of Physiology, 517:931–939, 1999.

[23] PM McAndrew, JB Dingwell, and JM Wilken. Walking variability during continuous pseudo-random oscillations of the support surface and visual field. Journal of Biomechanics, 43:1470–1475, 2010.

[24] E Anson, P Agada, T Kiemel, Y Ivanenko, F Lacquaniti, and J Jeka. Visual control of trunk translation and orientation during locomotion. Experimental Brain Research, 232:1941–1951, 2013.

[25] D Hamacher, F Herold, P Wiegel, D Hamacher, and L Schegga. Brain activity during walking: A systematic review. Neuroscience and Biobehavioral Reviews, 57:310–327, 2015.

[26] D Kindregan, L Gallagher, and J Gromley. Gait deviations in children with autism spectrum disorders: A review. Autism Research and Treatment, 2015, 2015.

[27] SM Morton and A J Bastian. Mechanisms of cerebellar gait ataxia. The Cerebellum, 6:79–86, 2007.

[28] T McGeer. Passive dynamic walking. The International Journal of Robotics Research, 9, 1990.

[29] S Collins, A Ruina, R Tedrake, and M Wisse. Efficient bipedal robots based on passive-dynamic walkers. Science, 307:1082–1085, 2005.

[30] K Byl and R Tedrake. Metastable walking machines. The International Journal of Robotics Research, 28:1040– 1064, 2009.

[31] Kevin C Galloway, Galen Clark Haynes, B Deniz Ilhan, Aaron M Johnson, Ryan Knopf, Goran Lynch, Benjamin Plotnick, Mackenzie White, and Daniel E Koditschek. X-rhex: A highly mobile hexapedal robot for sensorimotor tasks. University of Pennsylvania, Tech. Rep, 2010.

[32] GA Cavagna, NC Heglund, and CR Taylor. Mechanical work in terrestrial locomotion: two basic mechanisms for minimizing energy expenditure. American Journal of Physiology-Regulatory, Integrative and Comparative Physiology, 233:R243–R261, 1977.

[33] R Blickhan. The spring-mass model for running and hopping. Journal of Biomechanics, 22:1217–1227, 1989.

[34] RM Ghigliazza, R Altendorfer, P Holmes, and D Koditschek. A simply stabilized running model. SIAM Review, 43:519–549, 2005.

[35] A Seyfarth, H Geyer, M Gunther, and R Blickhan. A movement criterion for running. Journal of biomechanics, 35:649–655, 2002.

[36] JE Seipel and P Holmes. Running in three dimensions: Analysis of a point-mass sprung-leg model. The International Journal of Robotics Research, 24:657–674, 2005.

[37] JE Seipel and P Holmes. Three-dimensional translational dynamics and stability of multi-legged runners. The International Journal of Robotics Research, 25:889–902, 2006.

[38] AD Kuo. Stabilization of lateral motion in passive dynamic walking. The International Journal of Robotics Research, 18:917–930, 1999.

[39] MH Dickinson, CT Farley, RJ Full, MAR Koehl, R Kram, and S Lehman. How animals move : An integrative view. Science, 288:100–106, 2000.

[40] HJ Chiel and RB Beer. The brain has a body: adaptive behavior emerges from interactions of nervous system, body and environment. Trends in Neurosciences, 20:553–557, 1997.

[41] K Nishikawa, AA Biewener, P Aerts, AN Ahn, HJ Chiel, MA Daley, TL Daniel, RJ Full, MH Hale, TL Hedrick, AK Lappin, TR Nichols, RD Quinn, RA Satterlie, and B Szymik. Neuromechanics: An integrative approach for understanding motor control. Integrative and Comparative Biology, 47:16–54, 2007.

[42] HL More, JR Hutchinson, DF Collins, DJ Weber, SKH Aung, and JM Donelan. Scaling of sensorimotor control in terrestrial mammals. Proceedings of the Royal Society B: Biological Sciences, 277:3563–3568, 2010.

[43] HL More and JM Donelan. Scaling of sensorimotor delays in terrestrial mammals. Proceedings of the Royal Society B, 285:20180613, 2018.

[44] J Nirody. Flexible locomotion in complex environments: the influence of species, speed and sensory feedback on panarthropod inter-leg coordination. Journal of Experimental Biology, 226:jeb245111, 2023.

[45] MA Daley, G Felix, and AA Biewener. Running stability is enhanced by a proximo-distal gradient in joint neuromechanical control. Journal of Experimental Biology, 100:383–394, 2007.

[46] JJ Collins and IN Stewart. Coupled nonlinear oscillators and the symmetry of animal gaits. Journal of Nonlinear Science, 3:349–392, 1993.

[47] AS Machado, HG Marques, DF Duarte, DM Darmohray, and MR Carey. Shared and specific signatures of locomotor ataxia in mutant mice. eLife, page e55356, 2020.

[48] U Klibaite, M Kislin, JL Verpeut, S Bergeler, X Sun, JZ Shaevitz, and SS-H Wang. Deep phenotyping reveals movement phenotypes in mouse neurodevelopmental models. Molecular Autism, 13:12, 2022.

[49] P-P Vidal, L Degallaix, P Josset, J-P Gasc, and KE Cullen. Postural and locomotor control in normal and vestibular deficient mice. The Journal of Physiology, 559:625–638, 2004.

[50] L Ruder, A Takeoka, and S Arber. Long-distance spinal neurons ensure quadrupedal locomotor stability. Neuron, 92:1063–1078, 2016.

[51] HH Yang, LE Brezovec, LS Capdevilla, QX Vanderbeck, A Adachi, RS Mann, and RI Wilson. Fine-grained descending control of steering in walking drosophila. Cell, 187:6290–6308, 2024.

[52] S Rossignol, R Dubuc, and J-P Gossard. Dynamic sensorimotor interactions in locomotion. Physiological Reviews, 86:89–154, 2006.

[53] P Ramdya and AJ Ijspeert. The neuromechanics of animal locomotion: From biology to robotics and back. Science Robotics, 8, 2023.

[54] Y Blum, SW Lipfert, J Rummel, and A Seyfarth. Swing leg control in human running. Bioinspiration and Biomimetics, 5:026006, 2010.

[55] R Gal and F Libersat. New vistas on the initiation and maintenance of the insect motor behaviors revealed by specific lesions of the head ganglia. Journal of Comparative Physiology A, 192:1003–1020, 2006.

[56] R Zacarias, S Namiki, G M Card, ML Vasconcelos, and AM Moita. Speed dependent descending control of freezing behavior in drosophila melanogaster. Nature Communications, 8:3697, 2018.

[57] V Caggiano, R Leiras, H Goñi-Erro, D Masini, C Bellardita, J Bouvier, V Caldeira, G Fisone, and O Kiehn. Midbrain circuits that set locomotor speed and gait selection. Nature, 553:455–460, 2018.

[58] V Joshi and M Srinivasan. A controller for walking derived from how humans recover from perturbations. Journal of The Royal Society Interface, 16(157):20190027, 2019.

[59] NS Patil, JB Dingwell, and JP Cusumano. Correlations of pelvis state to foot placement do not imply within-step active control. Journal of Biomechanics, 97:109375, 2019.

[60] ME Morris, F Huxham, J McGinley, K Dodd, and R Iansek. The biomechanics and motor control of gait in parkinson disease. Clinical Biomechanics, 16:459–470, 2001.

[61] W Ilg, H Golla, P Thier, and MA Giese. Specific influences of cerebellar dysfunctions on gait. Brain, 130:786–798, 2007.

[62] KE Webster, JR Merory, and JE Wittwer. Gait variability in community dwelling adults with alzheimer disease. Alzheimer Disease and Associated Disorders, 20:37–40, 2006.

[63] SH Collins and AD Kuo. Two independent contributions to step variability during over-ground human walking. PloS One, 8:e73597, 2013.

[64] BL Rankin, SK Buffo, and JC Dean. A neuromechanical strategy for mediolateral foot placement in walking humans. Journal of Neurophysiology, 112(2):374–383, 2014.

[65] T Koolen, T de Boer, J Rebula, A Gowani, and J Pratt. Capturability-based analysis and control of legged locomotion, part 1: Theory and application to three simple gait models. The International Journal of Robotics Research, 31:1094–1113, 2012.

[66] ID Neveln, A Tirumalai, and S Sponberg. Information-based centralization of locomotion in animals and robots. Nature Communications, 10:3655, 2019.

[67] AI Goncalves, JA Zavatone-Veth, MR Carey, and DA Clark. Parallel locomotor control strategies in mice and flies. Current Opinion in Neurobiology, 73, 2022.

[68] BD DeAngelis, JA Zavatone-Veth, and DA Clark. The manifold structure of limb coordination in walking drosophila. eLife, page e46409, 2019.

[69] T Fujiwara, M Brotas, and ME Chiappe. Walking strides direct rapid and flexible recruitment of visual circuits for course control in drosophila. Neuron, 110:2124–2138, 2022.

[70] ME Chiappe. Circuits for self-motion estimation and walking control in drosophila. Current Opinion in Neurobiology, 81:102748, 2023.

[71] CJ Dakin, JT Inglis, R Chua, and J-S Blouin. Muscle-specific modulation of vestibular reflexes with increased locomotor velocity and cadence. Journal of neurophysiology, 110(1):86–94, 2013.

[72] M Bove, G Courtine, and M Schieppati. Neck muscle vibration and spatial orientation during stepping in place in humans. Journal of Neurophysiology, 88:2232–2241, 2001.

[73] YI Arshavsky, MB Berkinblit, OI Fukson, IM Gelfand, and GN Orlovsky. Origin of modulation in neurones of the ventral spinocerebellar tract during locomotion. Brain Research, 43:276–279, 1972.

[74] K Pfeiffer and U Homberg. Organization and functional roles of the central complex in the insect brain. Annual Review of Entomology, 59:165–184, 2014.

[75] A Mamiya, P Gurung, and JC Tuthill. Neural coding of leg proprioception in drosophila. Neuron, 100:636–650, 2018.

[76] BG Pratt, S-YG Lee, GM Chou, and JC Tuthill. Miniature linear and split-belt treadmills reveal mechanisms of adaptive motor control in walking drosophila. bioRxiv, 2024.

[77] A Santuz, T Akay, WP Mayer, TL Wells, A Schroll, and A Arampatzis. Modular organization of murine locomotor pattern in the presence and absence of sensory feedback from muscle spindles. The Journal of Physiology, 597:3147–3165, 2019.

[78] J Green, A Adachi, KK Shah, JD Hirokawa, PS Magani, and G Maimon. A neural circuit architecture for angular integration in drosophila. Nature, 546:101–106, 2017.

[79] AS Machado, DM Darmohray, J Fayad, HG Marques, and MR Carey. A quantitative framework for whole-body coordination reveals specific deficits in freely walking ataxic mice. eLife, page e07892, 2015.

[80] A Karayannidou, PV Zelenin, GN Orlovsky, and MG Sirota. Maintenance of lateral stability during standing and walking in the cat. Journal of Neurophysiology, 101:8–19, 2009.

[81] S Wang-Chen, VA Stimpfling, PG Özdil, L Genoud, F Hurtak, and P Ramdya. Neuromechfly 2.0, a framework for simulating embodied sensorimotor control in adult drosophila. bioRxiv, 2023.

[82] J Merel, D Aldarondo, J Marshall, Y Tassa, G Wayne, and B Ölveczky. Deep neuroethology of a virtual rodent. aRxiv, 2019.

[83] K Zakka, A Zeng, P Florence, J Tompson, J Bohg, and D Dwibedi. Xirl: Cross-embodiment inverse reinforcement learning. CORL, 2021.

[84] N Seethapathi, CB Clark, and M Srinivasan. Exploration-based learning of a stabilizing controller predicts locomotor adaptation. Nature Communications, 15:9498, 2024.

[85] Y Hurmuzlu and C Basdogan. On the measurement of dynamic stability of human locomotion. Journal of Biomechanical Engineering, 116:30–36], 1994.

[86] Christopher M Harris and Daniel M Wolpert. Signal-dependent noise determines motor planning. Nature, 394(6695):780–784, 1998.

[87] Sjoerd M Bruijn, OG Meijer, PJ Beek, and Jaap H van Dieen. Assessing the stability of human locomotion: a review of current measures. Journal of the Royal Society Interface, 10(83):20120999, 2013.

[88] J Englsberger, T Koolen, S Bertrand, J Pratt, C Ott, and A Albu-Schäffer. Trajectory generation for continuous leg forces during double support and heel-to-toe shift based on divergent component of motion. In IEEE/RSJ International Conference on Intelligent Robots and Systems, pages 4022–4019, 2014.

[89] M Shafiee, G Bellegarda, and A Ijspeert. Manyquadrupeds: Learning a single locomotion policy for diverse quadruped robots. International Conference of Robotics and Automation, 2024.

[90] SG Carver, NJ Cowan, and JM Guckenheimer. Lateral stability of the spring-mass hopper suggests a two-step control strategy for running. Chaos, 19, 2009.

[91] P Zaytsev, SJ Hasaneini, and A Ruina. Two steps is enough: No need to plan far ahead for walking balance. In IEEE International Conference of Robotics and Automation (ICRA), pages 6295–6300, 2015.

[92] MJP Toebes, MJM Hoozemans, R Furrer, J Dekker, and JH vanDieën. Local dynamic stability and variability of gait associated with fall history in elderly subjects. Gait and Posture, 36:527–531, 2012.

[93] A Nagamori, CM Laine, GE Loeb, and FJ Valero-Cuevas. Force variability is mostly not motor noise: Theoretical implications for motor control. PLoS Computational Biology, 17(3):e1008707, 2021.

[94] AS Voloshina and DP Ferris. Biomechanics and energetics of running on uneven terrain. Journal of Experimental Biology, 218:711–719, 2015.

[95] M Dhawale N, Venkadesan. How human runners regulate footsteps on uneven terrain. eLife, 12:e67177, 2023.

[96] MA Daley and AA Biewener. Running over rough terrain reveals limb control for intrinsic variability. Proceedings of the National Academy of Sciences, 103:15681, 2006.

[97] SR Datta, DJ Anderson, K Branson, P Perona, and A Leifer. Computational neuroethology: a call to action. Neuron, 104(1):11–24, 2019.

[98] EJ Dennis, A El Hady, A Michail, A Clemens, DR Gowan Tervo, J Voigts, and SR Datta. Systems neuroscience of natural behavior in rodents. Journal of Neuroscience, 41:911–919, 2021.

[99] A Mathis, P Mamidanna, KM Cury, Abe T, VN Murthy, MW Mathis, and M Bethge. Deeplabcut: markerless pose estimation for user-defined body parts. Nature Neuroscience, 21:1281–1289, 2018.

[100] TD Pereira, DE Aldarondo, L Willmore, M Kislin, SS-H Wang, M Murty, and JW Shaevitz. Fast animal pose estimation using deep neural networks. Nature Methods, 16:117–125, 2019.

[101] J Camargo, A Ramanathan, W Flanagan, and A Young. A comprehensive, open-source dataset of lower limb biomechanics in multiple conditions of stairs, ramps, and level-ground ambulation and transitions. Journal of Biomechanics, 112:110320, 2021.

[102] JJ Banks, W-R Chang, X Xu, and C-C Chang. Using horizontal heel displacement to identify heel strike in normal gait. Gait and Posture, 42:101–103, 2015.

[103] S Revzen and JM Guckenheimer. Estimating the phase of synchronized oscillators. Physical Review E, 78:051907, 2008.

[104] S Chatterjee. A new coefficient of correlation. Journal of the American Statistical Association, 116:2009–2022, 2020.

[105] Jr J.A Zeni, JG Richards, and JS Higginson. Two simple methods for determining gait events during treadmill and overground walking using kinematic data. Gait and Posture, 27:710,714, 2008.

[106] M Lemieux, N Josset, M Roussel, S Couraud, and F Bretzner. Speed-dependent modulation of the locomotor behavior in adult mice reveals attractor and transitional gaits. Frontiers in Neuroscience, 10, 2016.

[107] IE Marinescu, PN Lawlor, and KP Kording. Quasi-experimental causality in neuroscience and behavioural research. Nature Human Behaviour, 2:891–898, 2018.

